# The non-structural proteins NS3 and NS5A of Hepatitis C Virus (HCV) are degraded by two host proteolysis systems

**DOI:** 10.1101/2023.08.16.553615

**Authors:** Sara Mohamed, Jörg Soppa

## Abstract

Each lifecycle of the Hepatitis C virus (HCV) produces structural and non-structural (NS) proteins in equimolar. Structural proteins were either assembled or degraded by host proteolysis systems, while NS proteins remain inside the host cells and don’t accumulate. Therefore, they must be degraded. Here, NS3 and NS5A half-lives were quantified in the presence of autolysosome and proteasome different modulators. Inhibitors of both systems increased the half-life, while inducers decreased the half-life. Furthermore, polyubiquitination of NS3 and NS5A was observed. Additionally, their intracellular co-localization with autolysosome (LAMP2) and proteasome (PSMB5) was observed, and inhibitors of both systems increased the degree of co-localization. A better understanding of NS protein degradation might help to improve medical interventions during HCV infections in the future.

## Introduction

Chronic HCV infections lead to liver diseases and development of Hepatocellular carcinoma (HCC). About 71 million individuals are infected with HCV worldwide, in addition to 399,000 deaths occur due to HCV-associated liver diseases [2]. The direct-acting antiviral (DAA) therapy represents a breakthrough in HCV treatment, however there are still some limitations and complications of DAA therapy [1, 3, 45, 46]. Furthermore, HCV vaccine development has not been successful, underscoring the importance of efficient therapies [7, 8]. The goal of the WHO to overcome HCV infections by the year 2030 was interrupted by the COVID-19 pandemic, which led to a decrease in HCV diagnostics and treatment [9].

After infection, HCV RNA is translated into a single polyprotein, which is split into structural and non-structural (NS) proteins by viral and cellular proteases. Structural proteins form HCV viral particle, while NS proteins have various functions in HCV life-cycle [11, 16]. NS1 and NS2 are significant for viral assembly and release, while NS3-NS5B are forming the HCV replication complex (RC). Specifically, NS3/4A is involved in cleaving the 4-polyprotein junctions [12, 13], which highlights the significance of NS3 in HCV life-cycle. NS5B is an RNA-dependent RNA polymerase that is essential for HCV replication [14]. NS5A is a multifunctional protein, which is involved in HCV RNA replication and viral assembly, and it interacts with various different host proteins [15]. NS5A can be phosphorylated at multiple sites, hypophosphorylated and hyperphosphorylated variants, and have different roles in HCV lifecycle [33]. NS3, NS5A, and NS5B are the main targets for DAA therapy, based on their essential functions for HCV [45]. Therefore, we have focused on NS3 and NS5A.

HCV Structural and NS proteins are produced in equal amounts after polyprotein processing. Structural proteins are required in higher amounts than NS proteins, for the production of viral particles. However, it was observed that NS proteins don’t accumulate intracellularly, therefore, they must be degraded [14–16, 18]. Therefore, the aim of this study was to analyze whether NS proteins degradation occurs through one or both of the two major cellular degradation systems, the autophagosome and the proteasome.

Protein degradation via autophagosome and proteasome is very important to maintain cellular integrity [39]. Autophagy and proteasome are significant for HCV life-cycle [5, 19]. The close relationship between HCV and both degradation systems indicated that one or both systems are involved in NS proteins degradation. Previous reports have revealed different aspects of NS5A degradation [20–24]. In contrast, much less is known about NS3 degradation. Therefore, we used several biochemical and cell biological approaches to elucidate whether NS3 and NS5A are degraded via either or both degradation systems.

## Material and Methods

### Cell culture

The highly permissive human hepatic cell line derivative, Huh7.5, was used in this study [25]. Huh7.5 cells were grown in a complete media (CM) that was comprised of DMEM (Dulbecco’s modified Eagle’s medium; 4.5 g/I glucose; Sigma), 10% FCS (fetal calf serum), 0.1 U/ml penicillin, 100 μg/ml streptomycin and 2 mM L-glutamine. Cells were incubated in tissue culture flasks at 37°C, 5% CO_2_ and ∼90% humidity.

### Plasmids and in-vitro transcription

Plasmids pFK-JFH1/J6 (Jc1) and pJFH1/GND (GND) were described previously. GND was used as a replicative deficient variant [26, 27].

*In-vitro* transcription was performed using the linearized plasmids Cell script T7-Scribe^TM^ Standard RNA IVT kit (Biozym, Hessisch Oldendorf, DE) according to the manufacturer protocol. After *in vitro* transcription, Jc1 or GND RNA were electroporated to Huh7.5 cells as described before [18].

### Viability assay

Huh7.5 cells were seeded in a 96-well plate, then incubated with serial-dilutions of each modulator at 37°C. After 16 h treatment, the cells were incubated with 10% (w/v) PrestoBlue^TM^ cell viability reagent (ThermoFisher Scientific, Schwerte, DE) in CM. 1% (v/v) Triton-X 100 (Fluka, Deisenhofen, DE) was used as a positive control. Cells were monitored using a TECAN Reader (Infinite M1000, Tecan, Männedorf, Switzerland).

### Modulators of autophagy and proteasome

The Jc1 RNA was introduced into Huh7.5 cells by electroporation and then incubated for 56 h in CM. Subsequently, Huh7.5-Jc1 cells were incubated with different modulators (see below) for 16 h, while control cells were incubated for the same time in the absence of modulators. At 72 hours post electroporation (hpe), the effects of the modulators were quantified by comparison of treated and untreated cells using assays described below.

The following modulators were used:

100 µM of Leupeptin, 50 nM Bafilomycin-A1 (BFLA; Sigma Aldrich, Seelze, DE), 10 µM of SAR405 (Cayman, Bath, UK) were used to inhibit autophagy [28, 29, 47],

100 nM of Rapamycin (RAPA; Selleckchem, Boston, USA) was used to induce autophagy [47],

50 nM of Bortezomib (Selleckchem, Boston, USA) and 5 µM of PD169316 (Sigma Aldrich, Seelze, DE) was used to inhibit and induce proteasome, respectively [30, 48].

All inhibitors were dissolved in DMSO, except for Leupeptin, which is dissolved in water.

### Indirect Immunofluorescence Microscopy

At 72 hpe, Huh7.5-Jc1 cells were fixed with a mixture of Ethanol/Acetone (1:1), then immunofluorescence staining was performed using specific antibodies (1-2 h at RT). DAPI (4,6-Diamidin-2-phenylindol, Jackson ImmunoResearch Europe Ltd., Suffolk, UK) was used for nuclear staining. Mowiol (Sigma-Aldrich, Seelze, DE)-mounted cells were then analyzed with Leica SP8 CLSM (Leica, Wetzlar, Germany). The images were processed using the LASX software program.

Anti-NS3 (Mouse-monoclonal, Abcam, Cambridge, UK), anti-NS5A [31]; Rabbit-polyclonal), anti-PSMB5 (Rabbit-polyclonal, Abcam, Cambridge, UK, or Mouse-polyclonal, Sigma-Aldrich, Seelze, DE) and anti-LAMP2 (Guinea pig-polyclonal, Progen Biotechnik, Heidelberg, DE) were used as primary antibodies.

The following fluorophore-labeled secondary antibodies were used: anti-mouse/rabbit-IgG-Alexa 488, anti-mouse/rabbit-IgG –Flour-633 (Donkey polyclonal, ThermoFisher Scientific Schwerte DE), anti-mouse/rabbit-IgG-Cy3 (goat-polyclonal, Jackson ImmunoResearch Europe Ltd Suffolk UK).

To analyze protein-protein pixel co-localization, a tMOC (threshold Mander’s overlap coefficient) analysis was performed using the software Image Fiji [6, 10].

### SDS PAGE and Western Blot analysis

At 72 hpe, cells were lysed with RIPA buffer (50 mM Tris-HCl pH 7.2, 150 mM NaCl, 0.1% SDS (w/v), 1% Sodium desoxycholat (w/v) and 1% Triton X-100)). Then cells were sonicated, centrifuged to remove cellular debris, and proteins were quantified using Bradford assay.

Equal amounts of proteins, as detected by ß-actin protein, were loaded on SDS PAGE, then proteins were transferred to PVDF membranes. Membranes were then blocked either with 1% of Roti®-Block (Carl-Roth, Karlsruhe, DE) or with 10% (w/v) skim milk. For protein detection, membranes were incubated with specific antibodies (list below). The co-immunoperciptations were analyzed by the clean blot system (Thermo-scientific) to exclude any crossreativity with antibody fragments. A Li-COR Odessey CLx imaging system was used for protein detection and quantification of signal intensities. The results of the Co-IP experiments were analyzed using an Image Quant 800 imaging system.

Anti-GAPDH (anti-Glycerinaldehyd-3-phosphat-Dehydrogenase, Mouse monoclonal, Santa Cruz Biotechnology Incorporation, Dallas, USA), anti-Ubiquitin (Mouse monoclonal, In-vitrogen, Thermofischer Scientific), anti-NS5A, anti-NS3, anti-P62, anti-LAMP2, anti-ß-actin (Mouse monoclonal, Sigma-Aldrich, Seelze, DE) were used as primary antibodies.

Anti-mouse/anti-rabbit/anti-goat-IgG-IRDye680RD (Donkey polyclonal, LI-COR Biosciences GmbH Bad Homburg DE), anti-mouse/anti-rabbit-IgG-IRDye800CW (Donkey polyclonal, LI-COR Biosciences GmbH Bad Homburg DE), anti-mouse/-rabbit-IgG-HRP (Donkey polyclonal, GE Healthcare, Freiburg, DE) were used as secondary antibodies.

### Co-immunoprecipitation

Co-immunoperciptation was performed as described previously [4]. CO-IP was performed by 5 µg of anti-NS3, anti-NS5A, or anti-ubiquitin antibody.

### Cycloheximide (CHX) chase assay

To determine the half-lives of NS3 and NS5A, Huh7.5 cells were electroporated and incubated with modulators as described above. At 72 hpe, the supernatants were removed, and a solution of 142 µM CHX in CM was added. CM lacking CHX was added to control samples from time point zero, which were used to normalize all samples from subsequent time points. After incubation with CHX for various times from 15 minutes to 4 hours (time points see Results), the respective cells were lysed by the addition of RIPA buffer and the concentrations of NS3 and NS5A were quantified by Western blot analysis as described above. Three biological replicates were performed, and mean values of the signal intensities and their standard deviations were calculated for all time points. The mean values were used to calculate the half-lives of NS3 and NS5A under all conditions using a non-linear regression equation (one phase exponential decay, Y0 was set to 1, and the plateau was set to 0). NS3 and NS5A half-lives were calculated using GraphPad Prism 9.0.0 program (GraphPad, San Diego, USA).

### Statistical analysis

Three biological replicates were performed for all analyses. The results of the replicates were used to calculate mean values and ±SEM (standard errors of the means). Significance was analyzed by two-tailed unpaired Student’s *t* tests in Graph Prism 9.0.0 software. Statistical significance is presented in the figures as follows: * = P<0.05, ** = P<0.01, *** = P<0.001, and ns as non-significant.

## Results

### Determination of the half-lives of NS5A and NS3 and influence of modulators of the autophagosome and the proteasome activity

The half-lives of NS5A and NS3 were studied in the absence and the presence of different inhibitors as well as inducers of the autophagy and the proteasome. To this end, hepatoma cells (Huh7.5) were electroporated with viral Jc1 RNA and incubated for 56 h to allow the production of viral proteins. Huh7.5-Jc1 cells were then treated for 16 hours with different modulators of the two protein degradation systems. The following substances were used: leupeptin (100µM) as a late inhibitor of autophagy, SAR405 (10µM) as an early inhibitor of autophagy, rapamycin (100nM) as an autophagic flux inducer, bortezomib (50nM) as proteasome inhibitor, and PD169316 (5µM) as proteasome inducer. As a control, Huh7.5-Jc1 cells were harvested 72hpe without any treatment (WO).

NS5A and NS3 half-lives were determined by inhibiting translation with cycloheximide (CHX) and quantifying the amounts of the two proteins at 11 time points during the next four hours (CHX chase assay). The decreasing protein amounts at these time points were normalized to the amount at time point zero, i.e. in cells that had not been treated with CHX. **Figure 1A** shows representative Western blot analyses of one biological replicate (out of three). As expected, NS5A shows two bands, which represent the hyperphosphorylated (upper band) and the hypophosphorylated (lower band) form of the protein [33]. As a control, an antibody against the host housekeeping protein β-actin was used. We have quantified both bands of NS5A and the sum of both bands were normalized to the housekeeping protein, β-actin. The NS5A amounts at the 12 time points were quantified, average values of the three biological replicates and their standard deviations were calculated, and non-linear regression analyses were performed to generate regression graphs (**Figure 1B**). The graph of the untreated control (black line) is shown together with the graphs of the cultures that were treated with the five modulators (green lines). The half-lives that were determined by the non-linear regression analyses are shown in **Figure 1C**. The NS5A half-life (t1/2) was calculated in untreated cells (WO) to be about 2.43 h. Incubation with both autophagy inhibitors led to a drastic increase of the NS5A half-life, treatment with SAR405 (early inhibitor of autophagy) increased the half-life to 4.65 h, and treatment with leupeptin (late inhibitor of autophagy) led to an increase to 3.58 h. The influence of the autophagy inducer rapamycin was not as big, nevertheless, it decreased the half-life to 2.17 h. Notably, both proteasome modulators influence the half-life of NS5A, the proteasome inhibitor bortezomib increased the half-life to 3.59 h, while, in contrast, the proteasome inducer PD169316 led to a decrease to 1.77 h. Taken together, incubation with inhibitors of both systems increased the NS5A half-life, while incubation with inducers decreased the half-life, strongly indicating that both major protein degradation systems of the hepatoma host cells are involved in NS5A turnover.

**Figure 1.**
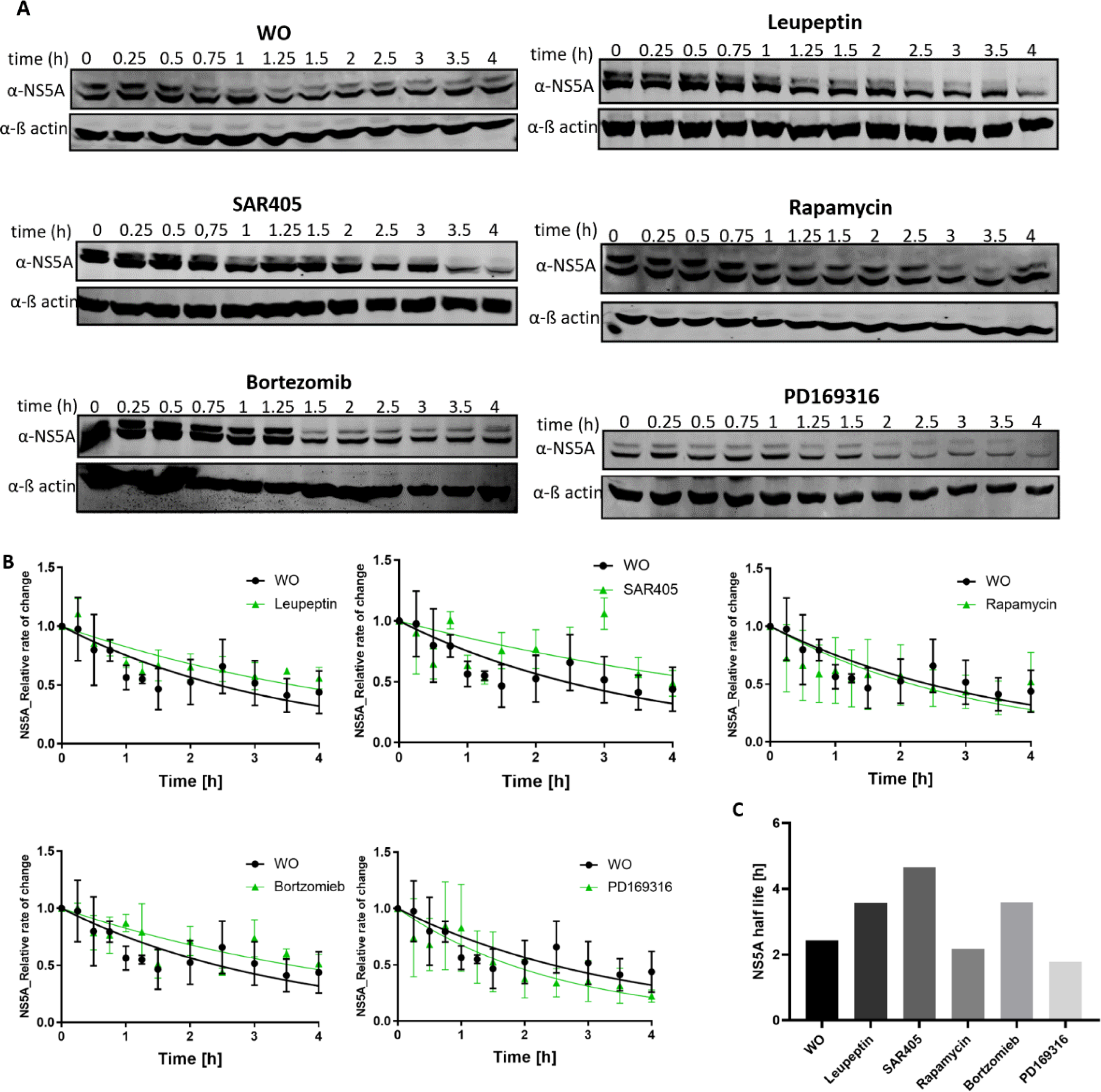
Determination of the NS5A half-life in the absence and presence of autophagy and proteasome modulators. **A**. The viral life cycle was started by electroporation of hepatoma cells with Jc1 RNA, and, after 56 h, the cultures were treated for 16 h with different modulators, as indicated, of left untreated. Translation was inhibited with CHX, and samples were taken at 11 different time points thereafter from 0.25 h to 4 h. Time point zero represents cells that were not treated with CHX. Representative Western blot analyses with an anti-NS5A antiserum and an anti-ß-actin as shown. **B.** The Western blot signals of three biological replicates were quantified, and average values and their standard deviations of the 12 time points are shown. In addition, non-linear regression graphs are shown. Black symbols/graphs represent the untreated control, and green symbols/graphs the cultures treated with the five indicated modulators. **C.** NS5A half-lives calculated from the regression graphs shown in B.

The same analyses were performed with an anti-NS3 antiserum, to determine the influence of the five modulators on the half-life of a second essential NS protein of HCV. **Figure 2A** shows representative Western blots from one biological replicate (out of three), **Figure 2B** shows the average values of the quantified Western blot signals and a non-linear regression graph, and **Figure 2C** shows the calculated half-lives of NS3. The NS3 half-life in the untreated control was about 24.2 h, much longer than the NS5A half-life of 2.43 h. It should be noted that due to its high stability only a small amount of NS3 was degraded during the 4 h experimental duration, therefore, the calculated NS3 half-lives are less exact than the NS5A half-lives as reported above. Nevertheless, the effects of the modulators were clearly visible. The two autophagy inhibitors, leupeptin and SAR405, considerably increased the half-life to calculated values of about 45.9 h and 44.8 h, respectively. In contrast, autophagic flux activation using rapamycin severely decreased the half-life to about 14.4 h. The Western blot signals after treatment with the proteasome inhibitor bortezomib did not follow the expected monophasic decay, therefore, the calculated value of about 50 h has to be taken with great care. In any case, the Western blot signals at least qualitatively revealed that NS3 was very stable in the presence of bortezomib. In stark contrast, induction of the proteasome with PD169316 drastically shortened the half-life to 8.54 h.

**Figure 2.**
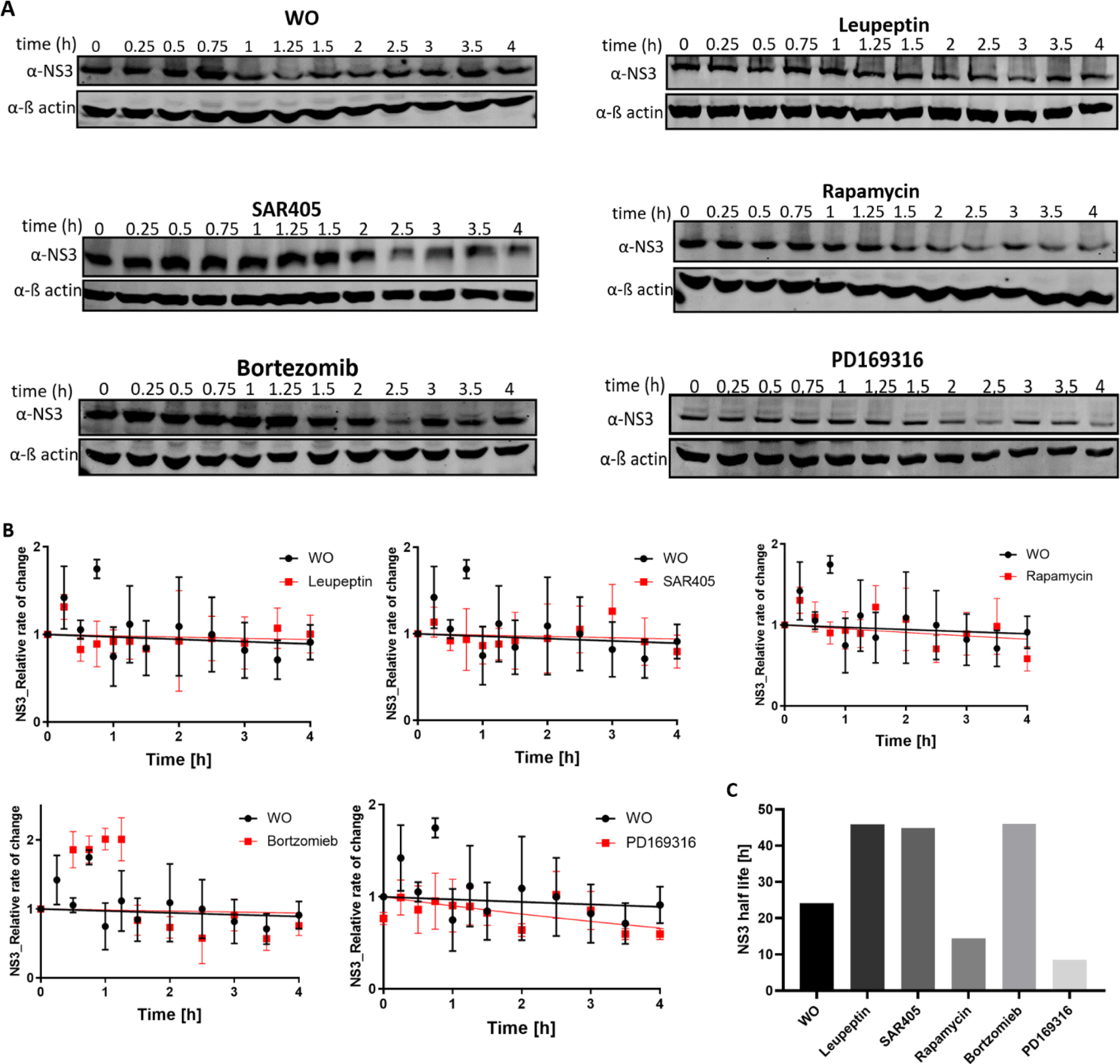
Determination of the NS3A half-life in the absence and presence of autophagy and proteasome modulators. **A**. The viral life cycle was started by electroporation of hepatoma cells with Jc1 RNA, and, after 56 h, the cultures were treated for 16 h with different modulators, as indicated, or left untreated. Translation was inhibited with CHX, and samples were taken at 11 different time points thereafter from 0.25 h to 4 h. Time point zero represents cells that were not treated with CHX. Representative Western blot analyses with an anti-NS3 antiserum and an anti-ß-actin as shown. **B.** Western blot signals of three biological replicates were quantified, and average values and their standard deviations of the 12 time points are shown. In addition, non-linear regression graphs are shown. Black symbols/graphs represent the untreated control, and green symbols/graphs the cultures treated with the five indicated modulators. **C.** NS3 half-lives calculated from the regression graphs shown in B.

Taken together, the combined results revealed that NS5A as well as NS3 were degraded by both protein degradation systems of the host, the autophagosome and the proteasome. In addition, the intracellular stabilities of the two proteins turned out to be very different, and NS3 had an about tenfold higher stability than NS5A.

### NS5A and NS3 poly-ubiquitination

The poly-ubiquitination of a protein is a cellular signal for protein degradation via the proteasome degradation system [30, 34]. To gain more insight into the proteasomal degradation of NS5A and NS3, co-immunoprecipitation (CO-IP) assays were performed in both directions. To this end, Huh7.5 cells were electroporated with Jc1 RNA to start the viral life cycle or with GND RNA (HCV non-replicating cells). At 72 hpe, cells were harvested and cytoplasmic extracts were generated. First, immunoprecipitation was performed with antisera against NS5A and against NS3, and Western blot analysis with an ubiquitin-specific antiserum was performed to reveal whether or not the two NS proteins were ubiquitinated. **Figure 3A** shows that indeed both proteins were polyubiquitinated. The size range was very large, indicating that the number of ubiquitin moieties (monomeric molecular weight 8.5 kDa) that are attached to both NS proteins is extremely variable and can be rather large. As expected, the cytoplasmic extracts (input) of both Huh7.5-Jc1 cells (NS protein producers) and Huh7.5-GND cells (non-producers) contained many polyubiquitinated proteins, showing that polyubiquitination is not restricted to viral NS proteins.

**Figure 3.**
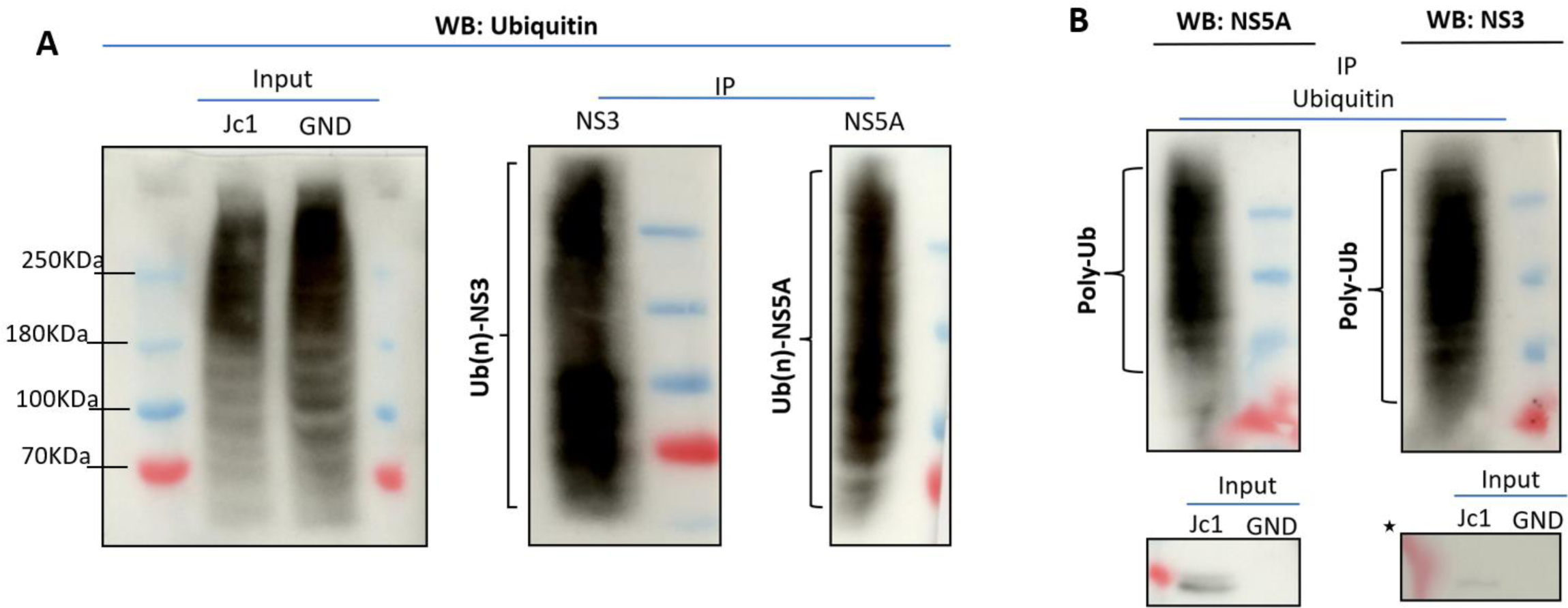
Poly-ubiquitination of NS5A and NS3. Huh7.5-Jc1 cells (HCV replicating) and Huh7.5-GND cells (negative control) were harvested 72hpe, and cytoplasmic extracts were prepared. Then co-immunoprecipitation analyses of NS5A and NS3 on the one hand and ubiquitin on the other hand were performed in both directions. **A.** Immunoprecipitation was performed with anti-NS5A and anti-NS3 antisera, and Western blot analysis of the precipitates and the input were performed with an anti-ubiquitin anti-serum. Representative Western blot membranes from one biological replicate are shown (out of three). **B.** Immunoprecipitation was performed with an anti-ubiquitin antiserum, and Western blot analysis of the precipitates and the input were performed with anti-NS5A and anti-NS3 antisera. Representative Western blot membranes from one biological replicate are shown (out of three). WB: Western Blotting, Ub(n)-: Ubiquitinated protein of interest, Poly-Ub: Poly-ubiquitin, the astrix mark * refer to an enhanced picture for better visualization.

Next, immunoprecipitation was performed with an anti-ubiquitin antiserum, and Western blot analysis was used with anti-NS5A and anti-NS3 antisera to reveal whether the two NS proteins were co-precipitated. **Figure 3B** shows that, indeed, Co-IP was successful in both directions. As expected, NS5A and NS3 could only be detected in the input (cytoplasmic extracts) from Huh7.5-Jc1 cells, but not in inputs from Huh7.5-GND (HCV non-replicating) cells, excluding that the Co-IP signals were due to cross-reactions of the two NS antisera to host proteins.

Taken together, Co-IP analyses in both directions revealed that NS5A as well as NS3 were polyubiquitinated, in agreement with their degradation via the proteasome shown above.

### Intracellular co-localization of NS5A and NS3 with the autophagosome and the proteasome

Fluorescence microscopy was used to study the intracellular co-localization of NS5A and NS3 with marker proteins of the autophagosome and the proteasome and the influence of modulators on the degree of co-localization. As before, the viral life cycle was started by electroporation of the hepatoma cell line Huh7.5 with Jc1 RNA and incubation of the cells for 56 hours. Then the modulators or DMSO were added, and the incubation was continued for 16 hours. At 72 hpe, cells were fixed, permeabilized, and treated with fluorescently-labelled antibodies. The marker proteins LAMP2 and PSMB5 were used to visualize the autophagosome and the proteasome, respectively. LAMP2 is a universal lysosome fusion marker with autophagosome in the final step of autolysosome formation [32]. PSMB5 is a protein from the inner ß subunits of the 20S core of the proteasome complex [34]. In addition, the DNA was stained with DAPI. Confocal Laser Scanning Microscopy (CLSM) was used for the intracellular localization of the antibody-decorated proteins. **Figure 4A** shows the co-localization of NS5A with the autophagosome. In this and the next Figures, the fluorescently labeled proteins are first shown separately, e.g. NS5A in the top left panel and LAMP2 in the middle left panel. Then, an overlay of the two pictures together with the DAPI stain is shown in the bottom left panel. In addition, a section of the overlay has been enlarged and is shown in the right panel. Pixel colocalizations of the different signals was quantified using tMOC (threshold Mander’s overlap coefficient) calculation analysis. The tMOC coefficient varies between 0 and 1, which equals 0% to 100% overlay. As can be seen in **Figure 4B**, the overlay between the NS5A and the LAMP2 signals was around 52% in DMSO treated cells. Treatment with Bafilomycin-A1 (a late inhibitor of autophagy) increased the overlay significantly to about 70%, while treatment with Rapamycin (an inducer of autophagy) reduced the overlay significantly to 33%.

**Figure 4.**
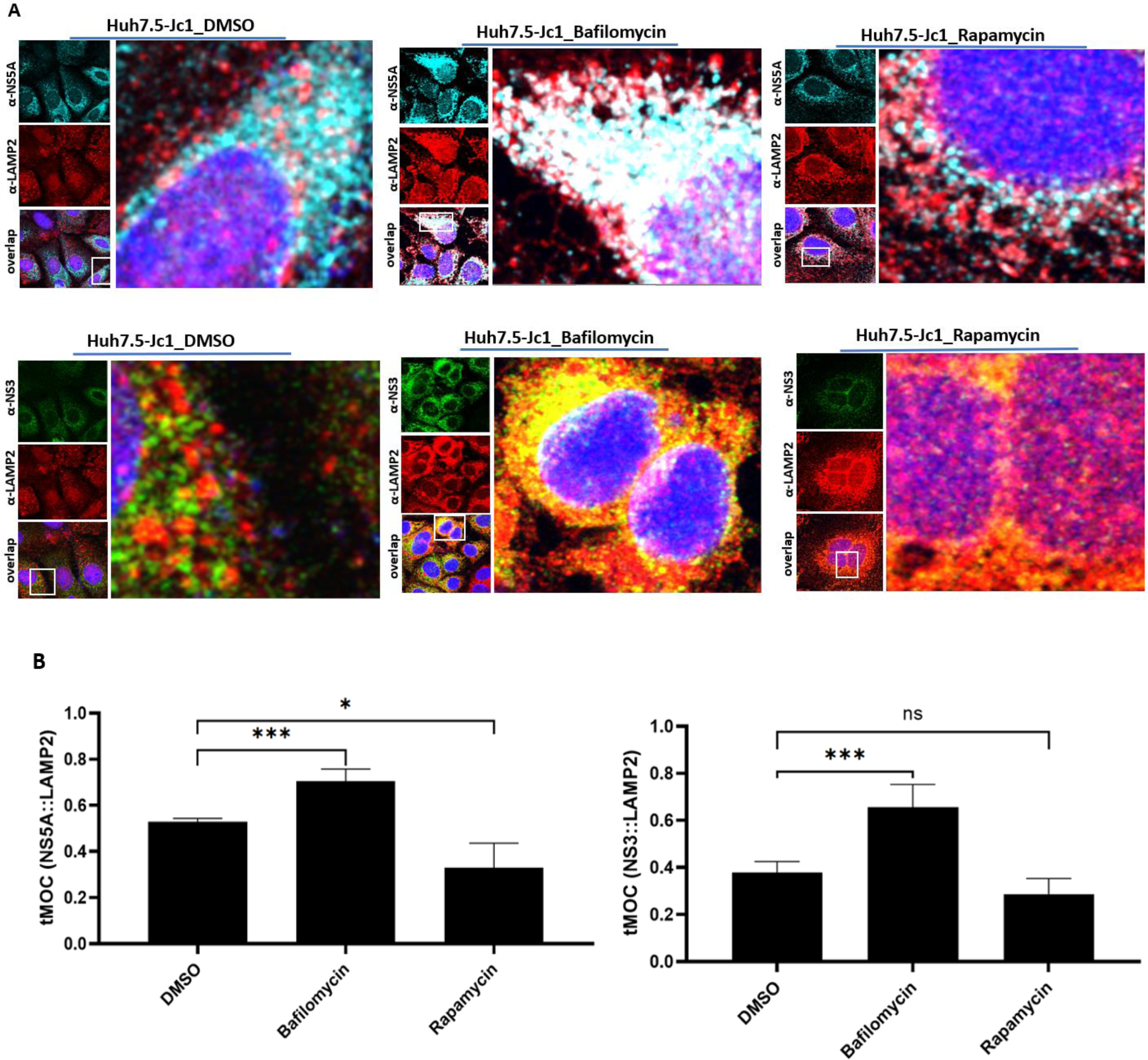
The intracellular co-localizations of NS5A and NS3 with the autophagosome. **A**. CLSM analysis of Huh7.5-Jc1 cells after treatment with different autophagy modulators and with DMSO, respectively. NS5A was stained with specific Alexa-633 (cyan blue; top left panel), NS3 with specific Alexa-488 (green; top left panel) and LAMP2 with cy3 (red; middle left panel). The bottom left panel shows an overlay of the two panels and a DAPI DNA-stain (blue). The right panel shows an enlargement of the area indicated in the overlay. Laser intensity and laser gain were kept constant. Magnification of 100X is shown. **B.** Quantification of the pixel co-localization of NS5A and NS3 with LAMP2 using a tMOC analysis. tMOC values were calculated from 6-8 cells of each replicate (n=3, mean ± standard error). The statistical significance was analyzed with a two-tailed unpaired *t*-test. * = p<0.05, *** = p<0.001, ns = non-significant.

Intracellular co-localization of NS3 with the autophagosome is shown in **Figure 4C**, and the tMOC quantification is shown in **Figure 4D**. The results are similar to the results with NS5A, i.e. the overlay is increased when the cells were treated with an inhibitor of autophagy, and it is decreased when the cells were treated with an inducer of autophagy. Therefore, in addition to the analyses described above, also the quantification of intracellular co-localization indicated that both NS proteins are degraded by the autophagy system of the host.

Next, intracellular co-localization of NS5A and NS3 with the proteasome was analyzed. **Figure 5A** shows co-localization of NS5A with the proteasome (PSMB5), and the result of the tMOC analysis is shown in **Figure 5B**. Treatment of the cells with the proteasome inhibitor bortezomib led to a significant increase of the overlap from 52% to about 66%. In stark contrast, treatment of the cells with PD169316 (induces the proteasome via inhibiting the p38 MAPK signaling pathway) [46] led to a severe decrease of the overlay to about 24%.

**Figure 5.**
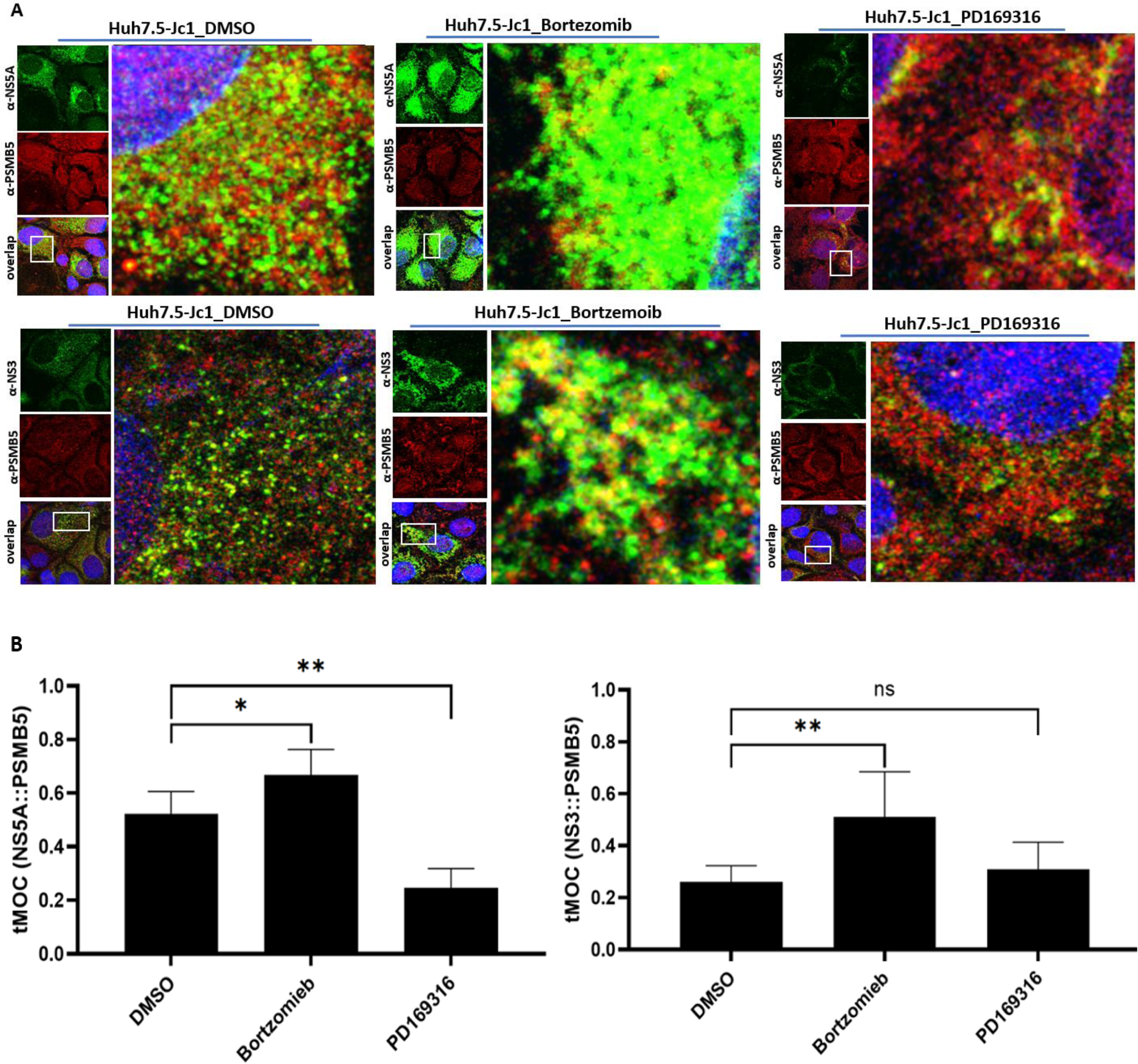
Intracellular co-localization of NS5A and NS3 with the proteasome. **A**. CLSM analysis of Huh7.5-Jc1 cells after treatment with different autophagy modulators and with DMSO, respectively. NS5A was stained with specific Alexa-488 (green; top left panel), NS3 with specific Alexa-488 (green; top left panel) and PSMB5 with cy3 (red; middle left panel). The bottom left panel shows an overlay of the two panels and a DAPI DNA-stain (blue). The right panel shows an enlargement of the area indicated in the overlay. Laser intensity and laser gain were kept constant. Magnification of 100X is shown. **B.** Quantification of the pixel co-localization of NS5A and NS3 with PSMB5 using a tMOC analysis. tMOC values were calculated from 6-8 cells of each replicate (n=3, mean ± standard error). The statistical significance was analyzed with a two-tailed unpaired *t*-test. * = p<0.05, ** = p<0.01, ns = non-significant.

The co-localization analysis of NS3 with the proteasome is shown in **Figure 5C**, and the tMOC quantification is shown in **Figure 5D**. Again, treatment of the cells with bortezomib led to a significant increase in the overlap from 26% to about 51%. However, in this case treatment of the cells with PD169316 did not result in a significant decrease of the overlap.

Notably, for both degradation systems the overlay was larger with NS5A (52%) than with NS3 (37 – 26%), indicating a higher degradation rate for NS5A. This is in excellent agreement with the shorter half-life of NS5A, compared to NS3 (see above). In summary, the quantification of the intracellular co-localization of NS5A and NS3 after treatment of virus-producing hepatoma cells with various modulators indicated that both NS proteins are degraded via both major protein-degradation systems of the host, the autophagosome and the proteasome.

## Discussion

During an HCV infection there are numerous interactions between viral and host functions. For example, HCV induces the formation of double membrane vesicles, which originate from the endoplasmic reticulum to perform the viral replication. Several reviews give an overview of HCV life cycle and how HCV proteins change functions of hepatoma host cell [35–37].

However, the present study focused on the opposite direction, the influence of host functions on viral proteins, specifically the degradation of two NS proteins by two host protein degradation pathways. It has been estimated that only 5% of NS proteins form the HCV RC [18]. Because the NS proteins don’t accumulate intracellularly, they must be degraded. Autophagy and ubiquitin-dependent proteasome are two major protein degradation pathways [39]. Therefore, we decided to study whether both of them are involved in the degradation of NS proteins. Earlier studies have already indicated their involvement; however, the results were partially contradictory (see below). Our aim was to resolve these contradictions and perform a study that is more systematic and exhaustive than prior studies. We concentrated on NS5A and NS3, because both are essential for the viral life-cycle, and if results apply to both proteins, it is likely that they can be generalized to additional NS proteins.

First, the half-lives of both NS proteins were determined in a hepatoma cell line infected with full-length HCV RNA. Previous studies have reported half-lives for NS5A, however, the values varied widely. The longest half-lives of 18.7 h and 16 h have been previously reported, however, shorter half-lives of 4-6 and 10 h were also reported [20, 40–42]. It was shown that after induction of proteasomal degradation by zinc mesoporphyrin, the half-life was drastically shortened from 18.7 h to 2.7 h [20], a value that is in excellent agreement with our calculated value of 2.4 h. Therefore, it seems that the experimental design of the present study, i.e. a 72 h incubation of cells after electroporation with a full-length HCV RNA, is a good mimic of an ongoing viral infection that includes active NS protein degradation. It should be noted that the levels of both isoforms of NS5A, hyperphosphorylated and hypophosphorylated variants, were used to determine the half-life (**Figure 1**). The half-lives of the two isoforms were not quantified, because the double bands were rather close together and it was not always easy to quantify them separately.

In two previous studies the NS3 protein half-life has been determined as 12 h and 14.5 h [17, 40]. In the present study a value of 24 h was calculated, which is somewhat larger than the values of previous studies. It should be noted that NS3 is severely stabilized by complex formation with NS4A, a cofactor for the serine protease activity of NS3. It has been shown that NS3 has a half-life of only 3 h when produced alone, but of 26 h when co-produced with NS4A [43].

Only one of the previous studies determined the half-lives of both proteins and found them to be nearly identical with 16 h and 14.5 h [40]. In stark contrast, we determined an about tenfold difference with 2.4 h for NS5A and 24 h for NS3. Several explanations for this difference seem possible, e.g. 1) we used a full-length HCV RNA, while the other study used a subgenomic fraction, 2) we used freshly transfected cells and determined the half-lives at 72 hpe, while the other study used continuous cultures in which the presence of the subgenomic HCV replicon was selected with Geneticin. Whatever the reason may be, the differences indicate that the experimental setup (e.g. cell lines, HCV constructs, cell culture, etc.) can have a large influence on the results.

Next, the influence of inhibitors and activators of both degradation systems on NS3 and NS5A half-lives was determined to elucidate whether one or both of these systems are involved in their turnover. Inhibitors of both degradation systems increased NS proteins half-lives, while activators of both systems decreased their half-lives. Therefore, the results unequivocally indicated that both NS proteins were degraded by both degradation systems. It has been reported before that NS3 is degraded by the proteasome [44]. However, it was proposed that an ubiquitination-independent pathway is used, while, in contrast, we found that both NS proteins are heavily polyubiquitinated.

A further approach to unravel whether NS proteins are degraded via both protein degradation systems was the quantification of the intracellular co-localization with autophagosome (LAMP2) and proteasome (PSMB5) marker proteins. Co-localization of NS5A with both marker proteins was higher than co-localization of NS3 with the marker proteins, which is in good agreement with the shorter half-life of NS5A as described above. Co-localization of both NS proteins increased when inhibitors of the autophagosome and the proteasome were applied, and it decreased when activators were applied. Therefore, also the co-localization approach indicated that both proteins were degraded by both systems.

Taken together, several experimental approaches revealed that the two essential HCV NS proteins, NS5A and NS3, are degraded by the autophagosome and the proteasome. Both proteins were polyubiquitinated, and, thus, proteasomal degradation seems to follow the classical pathway of proteasomal degradation, instead of the ubiquitin-independent pathway, as has been described above. NS5A and NS3 (as well as NS5B) are the main targets of DAA therapy. Therefore, knowledge that two protein degradation pathways are involved in their turnover might lead to further development of treatment strategies, e.g. by using combination therapies addressing both degradation pathways. A major aim would be to decrease the recurrence of HCV after DAA therapy. The development of DAA treatment has been a game changer in the fight against HCV [45]. However, recurrence of HCV-infection after successful treatment is a current problem [1, 45], and thus, any information on processes happening during HCV infection might help to solve this problem.

## Acknowledgements

The experimental part of the study was performed at the Paul-Ehrlich-Institute, D-63225 Langen, Germany. Sara Mohamed received a scholarship from the “Deutsche Akademischer Austauschdienst” (DAAD, German Academic Exchange Service).

